# The Nicotiana tabacum Mediator subunit MED25 regulates nicotine biosynthesis through interacting with the basic helix-loop-helix (bHLH) transcription factor NtMYC2s

**DOI:** 10.1101/2021.05.24.445359

**Authors:** Ge Bai, Yong Li, Da-Hai Yang, Tao Pang, Zhi-Yong Fan, Jia-En Shu, Mingliang Fei, Bingguang Xiao, Zhen-Yu Wang, He Xie

## Abstract

Nicotine is one of the most important secondary metabolites in tobacco, and its biosynthesis can be induced by topping and jasmonic acid treatment. NtMYC2s play pivotal roles in the regulation of nicotine. The mediator server as a bridge betwen the transcription factors and RNA polymerase in order to facilitates transcription and functions in plants. However, the role of mediator in the regulation of nicotine biosynthesis remains unknown. In this study, we firstly identify the NtMED25 through homologous analysis. NtMED25 interacts with NtMYC2s through the MD region. Interestingly, the nicotine content is decreased in the the knock-down transgenic lines of NtMED25, and the expression levels of two nicotine biosynthesis genes, NtQPT2 and NtPMT2, are also reduced when compared with that in the wild-type plants. Furthermore, NtMED25 enhances the binding of NtMYC2a/ NtMYC2b to the promoter of *NtPMT2* and *NtQPT2*, and then facilitates the nicotine biosynthesis. Therefore, our study revealed the function of mediator in the regulation of nicotine, and provide the insight role on the transcriptional regulation of plant secondary metabolites.

## Introduction

Tobacco (*Nicotiana tabacum*) can produce a variety of alkaloids that account for 0.6−3.0% of the dry matter mass in the leaves, which can protect them from insects feeding [1–3]. Among the alkaloids, the main ingredient is nicotine. The biosynthesis of nicotine is derived from the production of putrescine. Ornithine and arginine can be catalyzed to form putrescine through ornithine decarboxylase (ODC) or arginine decarboxylase (ADC), respectively [4]. S-adenosylmethionine (SAM) and putrescine are catalyzed by putrescine N-methyltransferase (PMT) into N-methyl putrescine (NMP) and S-adenosine homocysteine (SAH) [5]. NMP is oxidized by N-methtlputrescine oxidase (MPO) to produce 4-methylaminobutyraldehyde. The spontaneous cyclization of 4-methylaminobutyraldehyde can form the N-methyl-Δ’-pyrrolidine salts, which can be condensed with pyridine rings to produce nicotine [6]. The pyridine ring of nicotine is provided by nicotinic acid and the precursor of nicotinic acid is quinolinic acid that is produced from L-aspartic acid. The quinolinic acid is catalyzed by quinolinic acid phosphate ribosyltransferase (QPRT) to form the nicotinamide adenine dinucleotide (NAD), which is further cycled through the pyridine nucleotide pathway to nicotinic acid [7,8]. The biosynthesis of nicotine can be induced by herbivore feeding, insect biting, and pathogen attacking [9], as well as jasmonic acid and its derivatives (JAs), mechanical wounding and topping [10–12].

JAs are one of the most important phytohormones in planta. JAs can regulate the multiple processes of plant growth and development, including root growth, pollen development, fruit ripening, senescence and the biosynthesis of plant secondary metabolites [13–17]. It has also been found that JAs are participate in various biotic and abiotic stresses such as pathogen infection, insect feeding, mechanical wounding, salt stress[18–22]. To date, the pathway of JA-mediated nicotine synthesis has been well investigated. In the presence of JA, its derivative JA-Ile can bind to the JA receptor COI1 to result in ubiquitin degradation of the negative regulator JAZ protein, which can release the downstream transcriptional activators and activate jasmonate response in plants [23]. In tobacco, the regulators of the JA pathway COI1 and JAZ proteins have been demonstrated to be regulatory factors for the nicotine biosynthesis[11]. Moreover various transcription factors that can regulate the nicotine biosynthesis were identified including JAP1, ERF32 and ORC1 from ERF transcription factor family [24], bHLH1/2 and MYC2 from bHLH transcription factor family [25,26]. Furthermore, these ERF and bHLH transcription factors can affect the nicotine metabolism through their cross-regulation [12,27].

The signal pathway integration not only occurs at the level of transcriptional regulation and protein posttranslational modification such as phosphorylation and ubiquitination, but also includes the transcription initiation through the mediator complex and RNA Polymerase II[28,29]. The Pre-initiation complex (PIC) with RNA polymerase II as its core component is the functional unit to initiate transcription in the eukaryotes [29]. Numerous studies showed that mediator complex play an essential role in the regulation of signal transduction in eukaryotes. The mediator was firstly found and functionally investigated in yeast [28]. It was then found that human mediator plays a key role in multiple signaling pathways including Wnt-β-catenin [30], Hedgehog [31], RAS-MAPK [32] and TGFβ-SMAD [33]. Since the identification of mediator in the model plant *Arabidopsis Thaliana* [34], many studies have been carried out on the role of the mediator in the integration of plant signal pathways. *Arabidopsis* mediator AtMED25 is considered as an integrator of multiple signaling pathways and more than 20 proteins have been reported to interact with AtMED25 [35]. It has been found that AtMED25 mediates the transduction of JA signal through AtMYC2, which activates the downstream disease resistance process. Moreover, AtMED25 also interacts with AtABI5, an important transcription factor in the ABA pathway [36]. Meanwhile, AtMED16 can bind to EIN3 and EIL1 which are two key transcription factors involved in ethylene (ET) signaling pathways through AtMED25, and AtMED16 was positively regulating iron ion assimilation [37]. However, MED25 was involved in the regulation of nicotine biosynthesis remain unknown. In the current study, expressions of the nicotine biosynthesis genes *NtQPT2* and *NtPMT2* were significantly decreased in the NtMED25 interference lines, and then the nicotine content was greatly reduced in the NtMED25 interference lines when compared with that in the wild-type plants. Yeast two hybrid assays showed that NtMED25 binds to NtMYC2 through the MD region. Importantly, NtMED25 interacts with NtMYC2s which resulted in enhancing the transactivation ability of NtMYC2a and NtMYC2b to the downstream genes NTPMT2 and NTQPT2, and then regulating the nicotine biosynthesis.

## Results

### Identification of NtMED25

Based on the sequence similarity, a homolog of AtMED25 in tobacco was identified and was named NtMED25. The coding sequence of NtMED25 includes 526 bases and consists of 841 amino acids. The protein sequences of MED25 from tobacco, *Arabidopsis*, maize, rice and capsicum showed the higher similarities, suggesting that MED25 protein is evolutionarily conservative among the plants. According to the domains in AtMED25 protein, the MED25 proteins can be divided into four parts, including vWF-A, MD, ACID and GD. Compared with other MED25 proteins, NtMED25 contains 60 more amino acids in the vWF-A region (Figure 1A). The evolutionary analysis showed that MED25 proteins from monocots maize and rice were clustered together while MED25 proteins from the Solanaceae family tobacco and capsicum were clustered together. *Arabidopsis* MED25 is evolutionarily located between the monocots and the Solanaceae family (Figure 1B).

**Figure 1.**
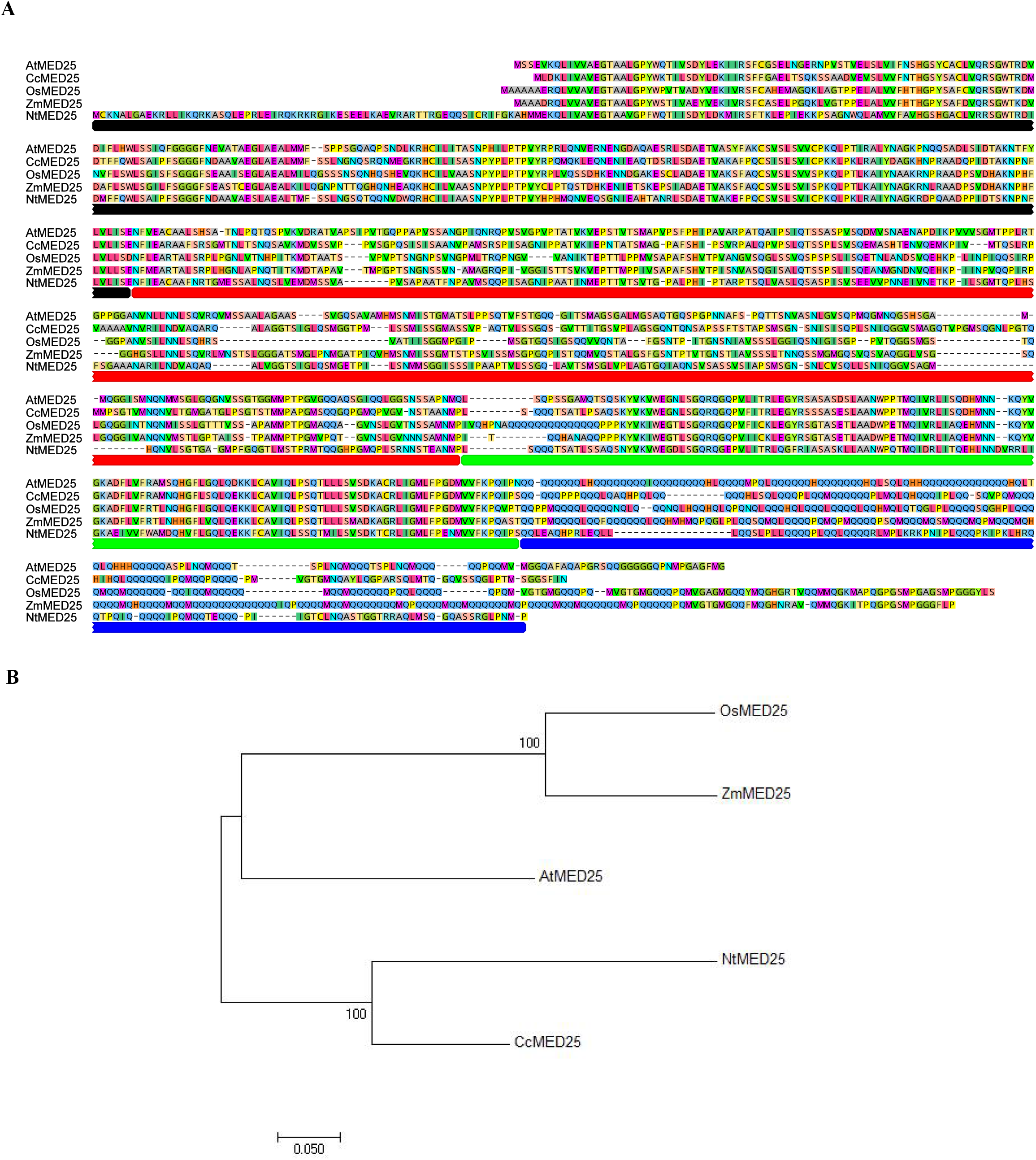
Idenfication of NtMED25. (A) The collinear analysis and domain analysis of MED25 proteins in tobacco, pepper, *Arabidopsis*, rice and corn. The red, yellow, blue and perple boxes indicate vWF-A domain, MD domain, ACID domain and GD domain, respectively. (B) The evolutionary tree of MED25 proteins from different species including Tobacco, pepper, *Arabidopsis*, rice and maize.

### NtMED25 Interacts with NtMYC2s in Tobacco

*Arabidopsis* MED25 physically interacts with the basic helix-loop helix transcription factor MYC2 though the ACID and the MD domain[30]. We therefore examined whether there was interaction between NtMED25 and NtMYC2 proteins (NtMYC2a and NtMYC2b) in tobacco. The NtMED25 sequences were cloned into pDEST32 vector, and the NtMYC2a and NtMYC2b sequences were cloned into pDEST22 vector, respectively. Similar to AtMED25, NtMED25 and NtMYC2b can activate the transcription of reporter genes in yeast after long-term culture, while there was none of self-activation after short-term culture. In accordance with *Arabidopsis* MED25, NtMED25 could interact with NtMYC2a or NtMYC2b in yeast, separately (Figure 2). To further elucidate the interaction domain between NtMED25 and NtMYC2, the NtMED25 protein was divided into four main regions, including NtMED25^vWF-A^, NtMED25^MD^, NtMED25^ACID^ and NtMED25^GD^. The sequences encoding of these four regions were cloned into pDEST32 vector, respectively, and then assay these possible interactions with NtMYC2a or NtMYC2b, respectively. Interestingly, only NtMED25^MD^ could interact with NtMYC2a as well as NtMYC2b. Moreover, the coding sequence of NtMED25^△vWF-A^, NtMED25 ^△GD^, NtMED25^MD-ACID^ and NtMED25^ACID-GD^ were cloned into pDEST32, respectively, and then map their interaction with NtMYC2a or NtMYC2b, respectively. The results showed that NtMED25^△vWF-A^, NtMED25 ^△GD^, NtMED25^MD-ACID^ could interact with NtMYC2a as well as NtMYC2b (Figure 2), indicating that MD domain is the critical for NtMED25 by interacting with NtMYC2 proteins.

**Figure 2.**
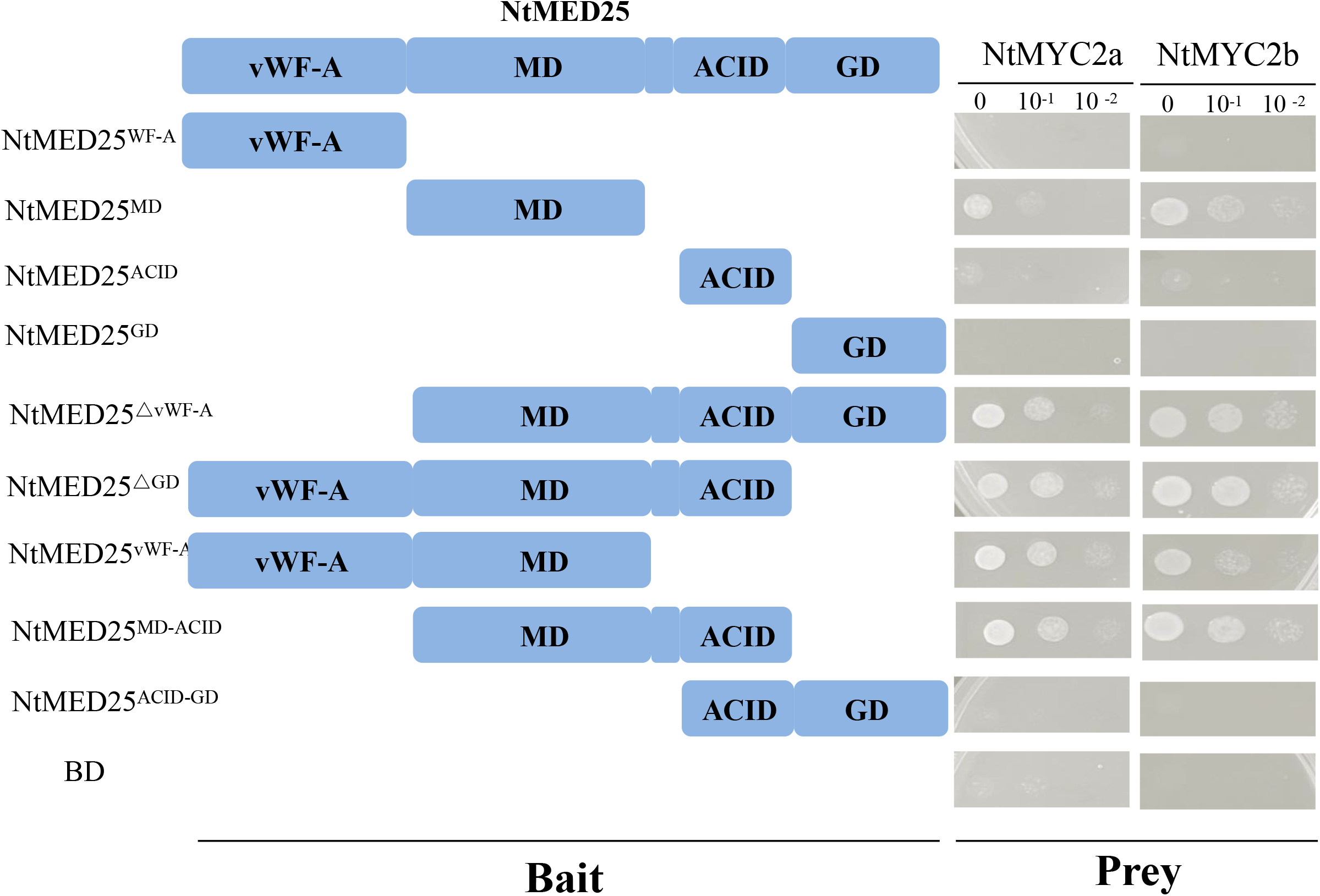
NtMED25 interacts with NtMYC2s in tobacco. Mapping of the domains involved in the NtMED25/NtMYC2 interaction using yeast two-hybrid assay. Based on the schematic protein structure of NtMED25 (top panel), full-length MED25 or its derivatives (PDEST22-NtMED25 or PDEST22-NtMED25 derivatives) were tested for interaction with MYC2 (PDEST32-MYC2) (see Methods for details). Transformed yeast was grown on selective media lacking His and Leu, and Trp (SD-3) to test protein interactions (left panel). The empty PDEST32 vector was cotransformed with MED25 and its derivatives in parallel as negative controls

### NtMED25 Affects the Nicotine Biosynthesis in Tobacco

AtMED25 is not only involved in the JA-induced resistance gene expression, but also participates in MYC2-mediated JA inhibition of taproot growth[31]. Meantime, NtMYC2 has been proved to be the core transcription factor in the nicotine biosynthesis that is mediated by JA in tobacco[26]. Therefore, we speculate that NtMED25 might be participated in the regulation of nicotine biosynthesis. We firstly obtained the knock-down transgenic lines of NtMED25 by RNA interference. It was showed that expression levels of *NtMED25* were significantly reduced in two transgenic lines by quantitative real-time PCR assay (Figure 3A). Expectedly, the nicotine content was much lower in two lines with reduction about 50% when compared with that in wild-type plants (Figure 3B). We then determine whether nicotine biosynthesis genes were differentiated expressed in response to JA. It was showed that the expressions of *NtPMT2* and *NtQPT2* were significantly induced under JA treatment in the wild-type plants (Figure 3C). Meanwhile, the transcript level of *NtPMT2* and *NtQPT2* were largely induced at 1 h after JA treatment and then decreased at 3 h and 6 h.In consist with nicotine content, the expressions of *NtPMT2* and *NtQPT2* were significantly lower in two transgenic lines than those in the wild-type plants (Figure 3C).

**Figure 3.**
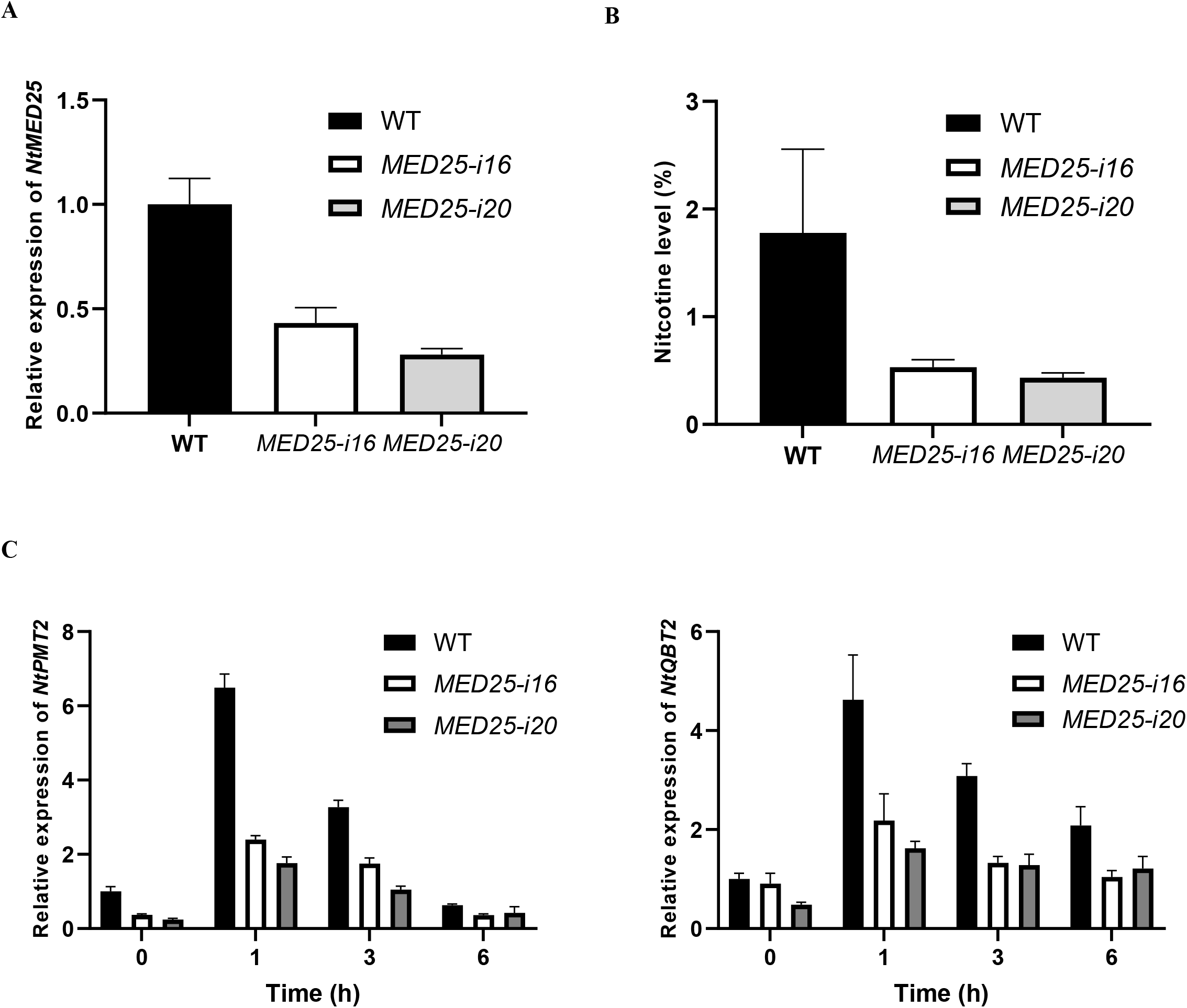
NtMED25 affected the nicotine content. (A) The expression levels of NtMED25 in two RNAi lines (*MED25-i16* and *MED25-i20*) were significantly decreased. (B) The nicotine contents in two RNAi lines were remarkably decreased. (C) The expression levels of NtQPT2 genes were decreased after JA treatment in two RNAi lines. (D) The expression levels of NtPMT2 genes were decreased after JA treatment in two RNAi lines.

### NtMED25 Enhances the Binding of NtMYC2a/ NtMYC2b to the Promoter of NtPMT2 and NtQPT2

Previous studies revealed that NtMYC2 proteins bind to the G-box motif in the NtPMT1a promoter to activate the nicotine biosynthesis in tobacco [26]. NtMED25 could interact with NtMYC2a/NtMYC2b (Figure 2), and NtMED25 plays pivotal role in regulating the nicotine biosynthesis in tobacco (Figure 3B). we therefore speculated that NtMED25 could serves as a bridge between NtMYC2 proteins and the RNA polymerase machinery to regulate the transcription of downstream genes. The reporter-effector transient expression assay system was performed in tobacco leaves. The promoter sequences of *NtQPT2* and *NtPMT2* genes were cloned into the vector pGreen0800 to form two reporter vector *Prob_QPT2_*::*LUC* and *Prob_PMT2_*::*LUC*, respectively. The coding sequences of *NtMYC2a*, *NtMYC2b* and *NtMED25* gens were cloned into the downstream of the 35S promoter to form activator vectors *Pro_35S_*:*NtMYC2a*, *Pro_35S_*:*NtMYC2b* and *Pro_35S_*:*NtMED25*, respectively (Figure 4A). The LUC assays showed that the expression levels of LUC were very low when only *Prob_QPT2_*:*LUC* or *Prob_PMT2_*:*LUC* was injected into tobacco leaves (Figure 4B). However, the fluorescence intensity of LUC was enhanced by 2-8 folds when *Prob_QPT2_*:*LUC* and *Prob_PMT2_*:*LUC* were injected into tobacco leaves together with *Pro_35S_*::*NtMYC2a* or *Pro_35S_*::*NtMYC2b* (Figure 4B). Furthermore, when *Pro_35S_*::*NtMED25* was injected into tobacco leaved together with *Pro_35S_*::*NtMYC2* and *Prob_QPT2_*::*LUC*/*Prob_PMT2_*::*LUC*, the fluorescence intensity of LUC was further increased. When *Pro_35S_*:*NtMED25* was injected, the LUC fluorescence intensity was twice as high as that of the absence of *Pro_35S_*:*NtMED25* (Figure 4B). These results suggesting that NtMED25 could promote the expression of *NtQPT2* and *NtPMT2* genes in cooperation with NtMYC2a and NtMYC2b.

**Figure 4.**
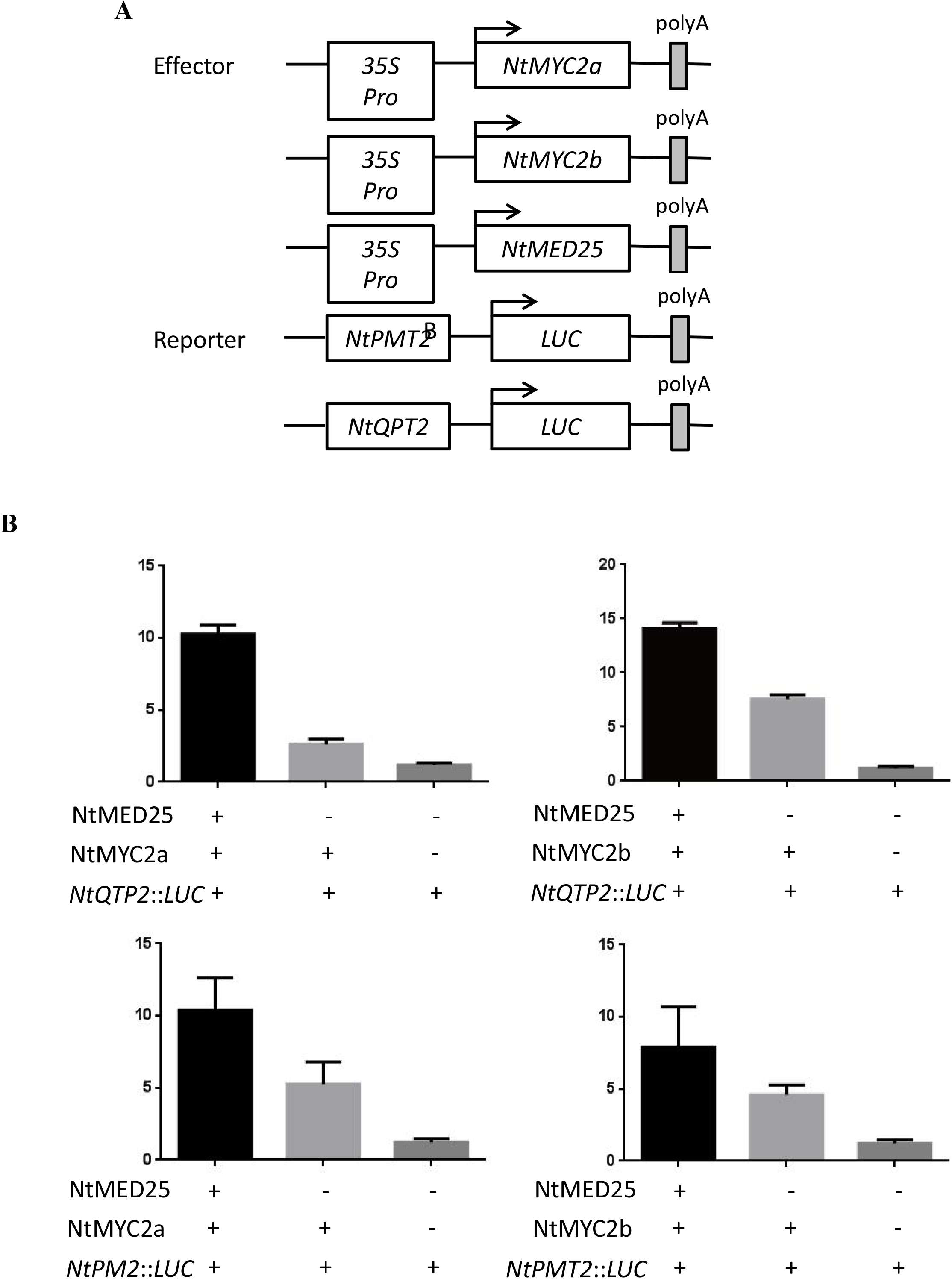
Dual-LUC analysis showed that NtMED25 enhanced the transcriptional activation of NtMYC2a and NtMYC2b to the promoters of NTQPT2 and NTPMT genes that are key enzymes in nicotine biosynthesis in tobacco. (A) The coding sequences of NtMED25, NtMYC2a and NtMYC2b were cloned into the overexpression vector, respectively, while the promoter sequences of *NtQPT2* and *NtPMT* genes were cloned into pGreen0800 vector. (B) The transcriptional activation of *NtQPT* and *NtPMT* was determined by comparing transformation of only MYC2a/MYC2b with transformation of MYC2a/MYC2b and NtMED25.

### The model of NtMED25 regulates NtMYC2-mediated dynamic transcriptional output of JA signaling

NtMED25 interacts with NtMYC2a/NtMYC2b, and then bridge the NtMYC2 proteins and the RNA polymerase machinery to regulate the transcription of *NtQPT2* and *NtPMT2* genes, resulted in the regulation of the nicotine biosynthesis in tobacco. Therefore, a model of NtMED25 involved in the transcriptional regulation of the nicotine biosynthesis was developed. In the absence of JA, JAZ proteins can interact with NtMYC2a and NtMYC2b to repress the transactivation ability of *NtMYC2* (Figure 5A), which results in lower expression levels of *NtPMT2* and *NtQPT2* genes that are the key enzyme genes in the nicotine biosynthetic pathway. Therefore, the lower content of nicotine was produced at this stage. However, when JA is present, JA promotes the binding of SCF^COI1^ to JAZ proteins, and JAZ is degraded through the 26S proteasome pathway (Figure 5B). NtMYC2a and NtMYC2b are further released from inhibition of JAZ proteins and promote their transactivation ability. Moreover, because NtMED25 can interact with NtMYC2a/NtMYC2b and recruit RNA polymerase machinery, the transactivation ability of NtMYC2a and NtMYC2b could be enhance in order to increase the transcript of *NTPMT2* and *NTQPT2* (Figure 5C), and then leads to the biosynthesis of nicotine.

**Figure 5.**
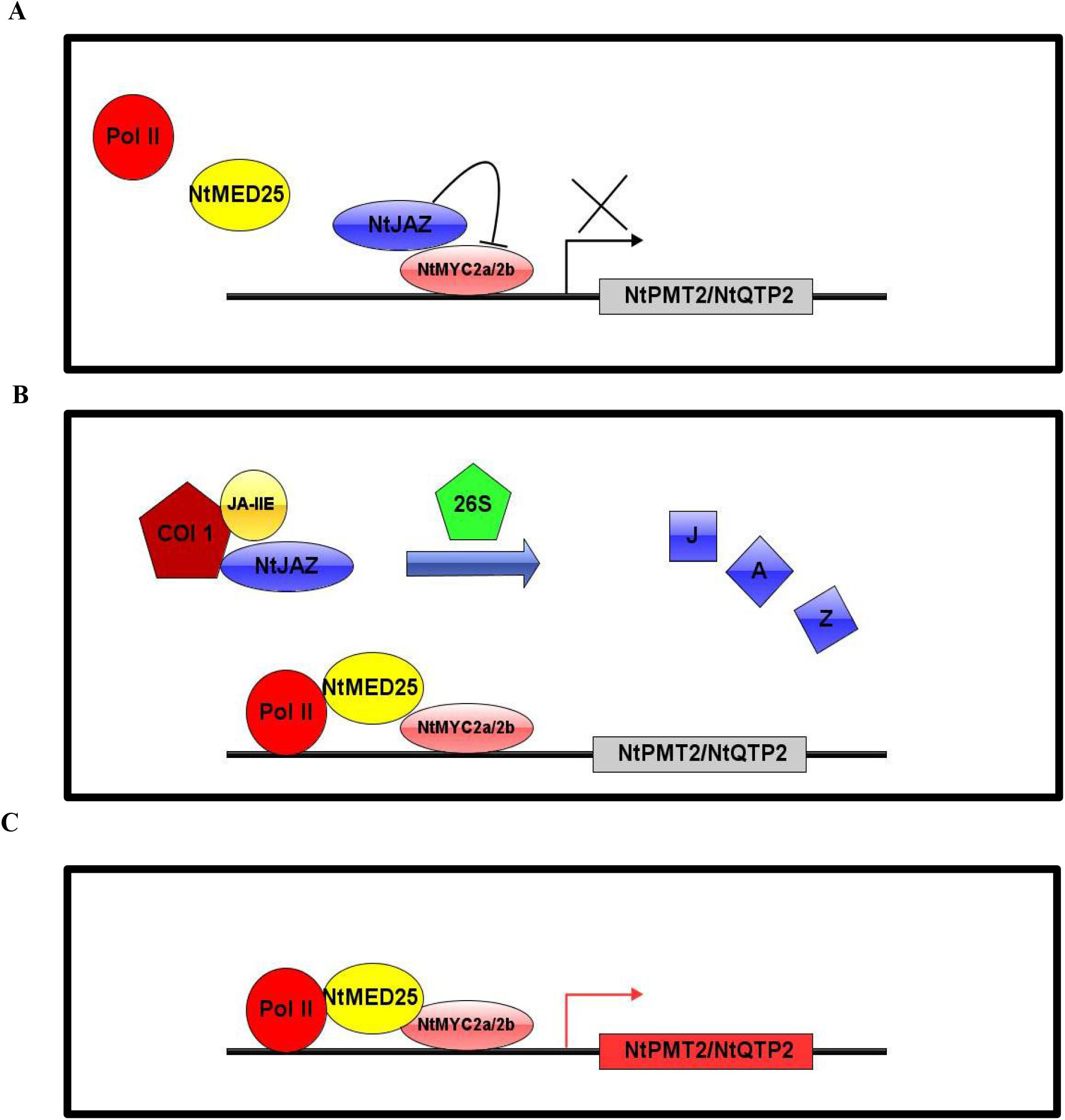
MED25 regulates MYC2-mediated dynamic transcriptional output of JA signaling. (A) In the absence of bioactive JA, JAZ proteins interact with MYC2 and repress MYC2 activity. The expression level of JA-responsive genes is low. (B) In the presence of JA-Ile, COI1 interaction with JAZ proteins, and subsequent degradation of JAZ repressors, which releases MYC2. (C) At the activation stage, MED25 physically interacts with Pol II and MYC2 form a transcriptional activator complex, which favors MYC2-mediated expression of JA-responsive genes. COI1, CORONATINEINSENSITIVE 1; JA-Ile, jasmonoyl-isoleucine; JAZ, jasmonate-ZIM domain; MED25, Mediator subunit 25; Pol II, RNA polymerase II.

## Discussion

The mediator plays an important role in the transcriptional regulation. The main function of the mediator is to connect the transcription factors with RNA polymerase machinery, which makes the transcription factors recruit RNA polymerase machinery and then transactivates the downstream genes. The function of mediator was firstly reported in animals and later investigated in plants. The main research currently focused on the development and stress resistance in plants [29–32]. However, the role of the mediator in plant secondary metabolism remains unknown.

Tobacco is one of the most important model plants, and nicotine is its most vital secondary metabolite. Nicotine has been deeply studied in tobacco, including the key enzyme genes and the transcriptional regulators involved in the biosynthesis pathway [26]. Among them, COI1-JAZ-MYC2 has been identified as the important transcriptional regulation pathway, which is induced by JA. It has been found that the expression of *NtMYC2* was increased when tobacco plants suffered topping, insect attacking or JA treatment [33]. COI1 bound to JAZ proteins, resulting in degradation of JAZ by 26S and release of NtMYC2 [11]. Thus, NtMYC2 could bind to the promoters of NtPMT2 and NtQPT2 genes, leading to their expression and finally the production of nicotine [26]. As a transcription factor, NtMYC2 needs to recruit Pol II, and there is a mediator to connect NtMYC2 and Pol II. According to the previous study, AtMED25, a tail protein of mediators in *Arabidopsis* can connect AtMYC2 to Pol II, which affects the expression of resistance genes *VSP1* and *PDF1.2* and the resistance to DC3000 [30]. Therefore, we speculated that the homolog of AtMED25 in tobacco could also function as a link protein between NtMYC2 and RNA polymerase machinery.

MED25 proteins from representative plants are evolutionally conservative, including dicotyledonous *Arabidopsis* and capsicum, and monocots corn and rice. All of them contain four domains including the vWF-A, MD, ACID and GD. The longer vWF-A domain in NtMED25 indicates their differentiated function. The polygenetic analysis showed that MED25 can be used to distinguish monocots from dicotyledons, and also distinguish cruciferous plants from solanaceous plants (Figure 1). The Y2H assays showed that NtMED25 could interact with NtMYC2a and NtMYC2b, which suggests that their interactions could facilitate NtMYC2a and NtMYC2b to activate the expression of *NtPMT2* and *NtQPT2* genes. The interactions between the different regions of NtMED25 and NtMYC2 showed that the NtMED25^MD^ was the key domain responsible for interaction between NtMED25 and NtMYC2 proteins (Figure 2). However, the key domain responsible for interaction between AtMED25 and AtMYC2 in *Arabidopsis* was AtMED25^MD-ACID^ and the two domains were indispensable. The key domains for interactions between MED25 and MYC2 proteins are different in *Arabidopsis* and tobacco, which suggests MED25 could function with the presence of the MED25^MD^ domain during the evolution.

The dual LUC assays were used to detect the transactivation ability in vivo using NtMED25, NtMYC2a and NtMYC2b as activators and the promoters of NtPMT2 and NtQPT2 genes as reporters. The expression of LUC reporter gene was almost not detected in *N. Benthamiana* leaves when only transformed with the reporter gene, suggesting that *NtQPT2* and *NtPMT2* genes were highly tissue-specific. They failed to be expression in leaves, which is the same as the previous results in tobacco [39,40]. The co-transformation of activators (NtMYC2a and NtMYC2b) and LUC reporters significantly enhanced the expression of LUC protein, suggesting that the expression of *NtPMT2* and *NtQPT2* genes were dependent on NtMYC2a and NtMYC2b. However, the results were different from those in BY2 cells with overexpression of *NtMYC2a* and *NtMYC2b* genes. The expression levels of *NtQPT2* and *NtPMT2* genes were not enhanced in BY2 cells with over-expression of NtMYC2a or NtMYC2b. However, the expression levels of *NtQPT2* and *NtPMT2* genes were decreased in the BY2 cells with NtMYC2 RNAi when JA treatment was applied [34]. The results in RNAi cells were consistent with our results that the expression of *NtQPT2* and *NtPMT2* genes depend on NtMYC2 genes. The inconsistent results in over-expression of NtMYC2a and NtMYC2b in BY2 cells may be due to the different plant materials. Importantly, the expression levels of *NtPMT2* and *NtQPT2* genes were significantly decreased before and after JA treatment in two NtMED25 knock-down transgenic lines, indicating that NtMED25 regulate the nicotine biosynthesis through differentiated expression of key enzyme genes.

Although it has been reported that mediators can facilitate transcription of the downstream genes, most of the researches were related to plant growth and development. It has been demonstrated that AtMED25 plays a role in promotion in expression of disease resistance genes in *Arabidopsis*. However, it was unclear whether MED25 has multiple functions. In the present study, we demostrated that NtMED25 could promote the expression of the key genes involved in the nicotine formation, suggesting NtMED25 plays a central role in the biosynthesis of plant secondary metabolites. Therefore, our results provide new insights into the functions of plant mediators and lay a basis for further research.

## Methods

### Plant Materials and Growth Conditions

The *Nicotiana tabacum L*. seeds were obtained from Yunnan Academy of Tobacco Agricultural Sciences (Yunnan, China). Surface-sterilized seeds were directly sowed into the soil in pots. The Nicotiana young seedlings were grown in the plant growth chamber with a 16-h-light/8-h-dark photoperiod under continuous white light (∼75 mol m−2 s−1) at 28 ℃-day/ 23 ℃-night. All plants were kept well-watered after sowing.

Tobacco samples were collected from plants in the field and flash-frozen in liquid nitrogen. Field management was performed according to regular agricultural practices.

### Phylogenetic and alignment

The protein sequences were obtained from China tobacco genome database V2.0 and NtMED25 proteins were predicted by HMMER [35]. NtMED25, AtMED25, ZmMED25, OSMED25 and CcMED25 were aligned with ClustalW software [36], and an unrooted phylogenetic tree was constructed using MEGA 7.0 (https://www.megasoftware.net/) with the neighbor-joining method and 1000 replicates of bootstrap [37].

### JA treatment

The 8-week-old tobacco plants were removed from pots and the roots were cleanly washed and immersed in water for 12 h. The roots were then immersed in 100 μM JA solution and were harvested after treatment for 1 h, 3 h and 6 h, respectively.

### RNA Extraction, cDNA Preparation and quantitative real time PCR

Total RNA was extracted with the SuperPure Plantpoly RNA Kit (GeneAnswer, BeiJing, China). All RNA samples were treated with RNase-free DNase I (GeneAnswer, BeiJing, China). NtMED25, NtPMT2, NtQPT genes were examined via qPCR and the method of qPCR were described by Bai et al.[38]. Breifly, A total of 2 μg of total RNA in a 20 μL reaction was converted to cDNA with a SuperScript III Reverse Transcriptase (Invitrogen, Waltham, Massachusetts, USA) by the manufacturer’s instructions on an Eppendorf Mastercycler thermocycler (Eppendorf AG, Germany). qPCR reactions were made with a SuperReal PreMix Plus SYBR Green Kit (TIANGEN Biotech, BeiJing, China) following manufacturer’s instructions in a 20 μL volume. qPCR was done on an Applied Biosystems™ QuantStudio™ 6 Flex Real-Time PCR System (ThemoFisher Scientific, Waltham, Massachusetts, USA). The log2fold change was calculated by the 2^-ΔΔCT^ method using 26S as a reference gene. The CT values represent the average of three technical replicates.

### Vector Construction

The RNAi fragment was amplified with NtMED25_RNAi_F and NtMED25_RNAi_R primers and the cDNA of N. Tabacum Yunyan87. The amplified fragment was cloned into pDONR/ZEO vector by the BP reaction with the Gateway BP Clonase II and further cloned into the pHellsgate12 vector through the LR reaction with Gateway LR Clonase. The Yunyan 87 plant leaves were used to transformation to produce RNAi plants according to the previous report. The different fragments of NtMED25 were cloned into pDEST22 vector while the sequences of NtMYC2a and NtMYC2b were cloned into Pdest32 vector, respectively.

#### Yeast two hybrids

Using the LiOAC method, each pair of AD and BD constructs were co-transformed into Saccharomyces cerevisiae strain AH109. were used for co-transformation with empty pDEST22 and pDEST32 vectors, respectively. Transforms were plated on the SD minimal media without leucine (Leu) and tryptophan (Trp). The positive yeasts were grown in liquid SD-Leu/Trp medium at 30 C for 1 day and 10 ll of dilutions of 1:10, 1:100, 1:1,000 and 1:10,000 were pipetted onto various plates (SD-Leu/Trp, SD-Leu/Trp/His)

## Author Contributions

Conceptualization, He Xie, Ge Bai; Data curation, Ge Bai, Yong Li; Formal analysis, Dahai Yang; Funding acquisition, Ge Bai, He Xie, Da-Hai Yang; Investigation, Ge Bai, Ming Liang Fei and Bingguang Xiao; Methodology, Zhi-Yong Fan, Jia-En Shu; Project administration, He Xie; Resources, MingLiang Fei, Bingguang Xiao, Methodology; Software Tao Pang; Supervision, Jun Yang; Writing – original draft, Ge Bai and He Xie; Writing – review & editing, Zhenyu Wang.

## Reference

1. Kessler, A.; Baldwin, I.T. Plant responses to insect herbivory: the emerging molecular analysis. Annual review of plant biology 2002, 53, 299–328, doi:10.1146/annurev.arplant.53.100301.135207.

2. Steppuhn, A.; Gase, K.; Krock, B.; Halitschke, R.; Baldwin, I.T. Nicotine’s defensive function in nature. PLoS biology 2004, 2, E217, doi:10.1371/journal.pbio.0020217.

3. Reddy, G.V.; Guerrero, A. Interactions of insect pheromones and plant semiochemicals. Trends in plant science 2004, 9, 253–261, doi:10.1016/j.tplants.2004.03.009.

4. Kakkar, R.K.; Sawhney, V.K. Polyamine research in plants–a changing perspective. Physiologia Plantarum 2002, 116, 281–292.

5. Biastoff, S.; Brandt, W.; Drager, B. Putrescine N-methyltransferase--the start for alkaloids. Phytochemistry 2009, 70, 1708–1718, doi:10.1016/j.phytochem.2009.06.012.

6. Katoh, A.; Shoji, T.; Hashimoto, T. Molecular cloning of N-methylputrescine oxidase from tobacco. Plant & cell physiology 2007, 48, 550–554, doi:10.1093/pcp/pcm018.

7. Chattopadhyay, M.K.; Ghosh, B. Molecular analysis of polyamine biosynthesis in higher plants. Current Science 1998, 517–522.

8. Wagner, R.; Wagner, K.G. The pyridine-nucleotide cycle in tobacco Enzyme activities for the de-novo synthesis of NAD. Planta 1985, 165, 532–537, doi:10.1007/BF00398100.

9. Baldwin, I.T. Functional interactions in the use of direct and indirect defences in native Nicotiana plants. Novartis Foundation symposium 1999, 223, 74-87; discussion 87-94, 160–165, doi:10.1002/9780470515679.ch6.

10. Baldwin, I.T.; Zhang, Z.-P.; Diab, N.; Ohnmeiss, T.E.; McCloud, E.S.; Lynds, G.Y.; Schmelz, E.A. Quantification, correlations and manipulations of wound-induced changes in jasmonic acid and nicotine in Nicotiana sylvestris. Planta 1997, 201, 397–404.

11. Shoji, T.; Ogawa, T.; Hashimoto, T. Jasmonate-induced nicotine formation in tobacco is mediated by tobacco COI1 and JAZ genes. Plant and cell physiology 2008, 49, 1003–1012.

12. Shoji, T.; Hashimoto, T. Tobacco MYC2 regulates jasmonate-inducible nicotine biosynthesis genes directly and by way of the NIC2-locus ERF genes. Plant and Cell Physiology 2011, 52, 1117–1130.

13. Feys, B.; Benedetti, C.E.; Penfold, C.N.; Turner, J.G. Arabidopsis Mutants Selected for Resistance to the Phytotoxin Coronatine Are Male Sterile, Insensitive to Methyl Jasmonate, and Resistant to a Bacterial Pathogen. The Plant cell 1994, 6, 751–759, doi:10.1105/tpc.6.5.751.

14. McConn, M.; Browse, J. The Critical Requirement for Linolenic Acid Is Pollen Development, Not Photosynthesis, in an Arabidopsis Mutant. The Plant cell 1996, 8, 403–416, doi:10.1105/tpc.8.3.403.

15. Falkenstein, E.; Groth, B.; Mithofer, A.; Weiler, E.W. Methyljasmonate and alpha-linolenic acid are potent inducers of tendril coiling. Planta 1991, 185, 316–322, doi:10.1007/BF00201050.

16. Wasternack, C.; Strnad, M. Jasmonates are signals in the biosynthesis of secondary metabolites - Pathways, transcription factors and applied aspects – A brief review. New biotechnology 2019, 48, 1–11, doi:10.1016/j.nbt.2017.09.007.

17. Chen, H.; Jones, A.D.; Howe, G.A. Constitutive activation of the jasmonate signaling pathway enhances the production of secondary metabolites in tomato. FEBS letters 2006, 580, 2540–2546, doi:10.1016/j.febslet.2006.03.070.

18. Browse, J. Jasmonate: an oxylipin signal with many roles in plants. Vitamins and hormones 2005, 72, 431–456, doi:10.1016/S0083-6729(05)72012-4.

19. Browse, J. Jasmonate passes muster: a receptor and targets for the defense hormone. Annual review of plant biology 2009, 60, 183–205, doi:10.1146/annurev.arplant.043008.092007.

20. Rao, M.V.; Lee, H.; Creelman, R.A.; Mullet, J.E.; Davis, K.R. Jasmonic acid signaling modulates ozone-induced hypersensitive cell death. The Plant cell 2000, 12, 1633–1646, doi:10.1105/tpc.12.9.1633.

21. Turner, J.G.; Ellis, C.; Devoto, A. The jasmonate signal pathway. The Plant cell 2002, 14 Suppl, S153–164, doi:10.1105/tpc.000679.

22. Wasternack, C. Jasmonates: an update on biosynthesis, signal transduction and action in plant stress response, growth and development. Annals of botany 2007, 100, 681–697, doi:10.1093/aob/mcm079.

23. Thines, B.; Katsir, L.; Melotto, M.; Niu, Y.; Mandaokar, A.; Liu, G.; Nomura, K.; He, S.Y.; Howe, G.A.; Browse, J. JAZ repressor proteins are targets of the SCF(COI1) complex during jasmonate signalling. Nature 2007, 448, 661–665, doi:10.1038/nature05960.

24. De Sutter, V.; Vanderhaeghen, R.; Tilleman, S.; Lammertyn, F.; Vanhoutte, I.; Karimi, M.; Inze, D.; Goossens, A.; Hilson, P. Exploration of jasmonate signalling via automated and standardized transient expression assays in tobacco cells. The Plant journal: for cell and molecular biology 2005, 44, 1065–1076, doi:10.1111/j.1365-313X.2005.02586.x.

25. Todd, A.T.; Liu, E.; Polvi, S.L.; Pammett, R.T.; Page, J.E. A functional genomics screen identifies diverse transcription factors that regulate alkaloid biosynthesis in Nicotiana benthamiana. The Plant Journal 2010, 62, 589–600.

26. Zhang, H.B.; Bokowiec, M.T.; Rushton, P.J.; Han, S.C.; Timko, M.P. Tobacco transcription factors NtMYC2a and NtMYC2b form nuclear complexes with the NtJAZ1 repressor and regulate multiple jasmonate-inducible steps in nicotine biosynthesis. Molecular plant 2012, 5, 73–84, doi:10.1093/mp/ssr056.

27. De Boer, K.; Tilleman, S.; Pauwels, L.; Vanden Bossche, R.; De Sutter, V.; Vanderhaeghen, R.; Hilson, P.; Hamill, J.D.; Goossens, A. APETALA2/ETHYLENE RESPONSE FACTOR and basic helix-loop-helix tobacco transcription factors cooperatively mediate jasmonate-elicited nicotine biosynthesis. The Plant journal: for cell and molecular biology 2011, 66, 1053–1065, doi:10.1111/j.1365-313X.2011.04566.x.

28. Kelleher III, R.J.; Flanagan, P.M.; Kornberg, R.D. A novel mediator between activator proteins and the RNA polymerase II transcription apparatus. Cell 1990, 61, 1209–1215.

29. Buendia-Monreal, M.; Gillmor, C.S. Mediator: A key regulator of plant development. Developmental biology 2016, 419, 7–18, doi:10.1016/j.ydbio.2016.06.009.

30. Chen, R.; Jiang, H.; Li, L.; Zhai, Q.; Qi, L.; Zhou, W.; Liu, X.; Li, H.; Zheng, W.; Sun, J. The Arabidopsis mediator subunit MED25 differentially regulates jasmonate and abscisic acid signaling through interacting with the MYC2 and ABI5 transcription factors. The Plant cell 2012, 24, 2898–2916.

31. Muñoz‐Parra, E.; Pelagio‐Flores, R.; Raya‐González, J.; Salmerón‐Barrera, G.; Ruiz‐Herrera, L.F.; Valencia‐Cantero, E.; López‐Bucio, J. Plant–plant interactions influence developmental phase transitions, grain productivity and root system architecture in Arabidopsis via auxin and PFT1/MED25 signalling. Plant, cell & environment 2017, 40, 1887–1899.

32. Elfving, N.; Davoine, C.; Benlloch, R.; Blomberg, J.; Brannstrom, K.; Muller, D.; Nilsson, A.; Ulfstedt, M.; Ronne, H.; Wingsle, G., et al. The Arabidopsis thaliana Med25 mediator subunit integrates environmental cues to control plant development. Proceedings of the National Academy of Sciences of the United States of America 2011, 108, 8245–8250, doi:10.1073/pnas.1002981108.

33. Li, F.; Zhang, H.; Wang, S.; Xiao, W.; Ding, C.; Liu, W.; Guo, H. Identification of Topping Responsive Proteins in Tobacco Roots. Frontiers in plant science 2016, 7, 582, doi:10.3389/fpls.2016.00582.

34. Yang, Y.; Guo, J.; Yan, P.; Li, Y.; Liu, K.; Gao, P.; Zhao, H.; Chen, Y.; Wang, Y.; Timko, M.P. Transcriptome profiling identified multiple jasmonate ZIM-domain proteins involved in the regulation of alkaloid biosynthesis in tobacco BY-2 cells. Plant molecular biology reporter 2015, 33, 153–166.

35. Finn, R.D.; Clements, J.; Arndt, W.; Miller, B.L.; Wheeler, T.J.; Schreiber, F.; Bateman, A.; Eddy, S.R. HMMER web server: 2015 update. Nucleic acids research 2015, 43, W30–38, doi:10.1093/nar/gkv397.

36. McWilliam, H.; Li, W.; Uludag, M.; Squizzato, S.; Park, Y.M.; Buso, N.; Cowley, A.P.; Lopez, R. Analysis Tool Web Services from the EMBL-EBI. Nucleic acids research 2013, 41, W597–600, doi:10.1093/nar/gkt376.

37. Kumar, S.; Stecher, G.; Tamura, K. MEGA7: Molecular Evolutionary Genetics Analysis Version 7.0 for Bigger Datasets. Molecular biology and evolution 2016, 33, 1870–1874, doi:10.1093/molbev/msw054.

38. Bai, G.; Yang, D.H.; Cao, P.; Yao, H.; Zhang, Y.; Chen, X.; Xiao, B.; Li, F.; Wang, Z.Y.; Yang, J., et al. Genome-Wide Identification, Gene Structure and Expression Analysis of the MADS-Box Gene Family Indicate Their Function in the Development of Tobacco (Nicotiana tabacum L.). International journal of molecular sciences 2019, 20, doi:10.3390/ijms20205043.

